# Frequency and synchrony of actomyosin oscillation during PCP-dependent convergent extension

**DOI:** 10.1101/316745

**Authors:** Asako Shindo, Yasuhiro Inoue, Makoto Kinoshita, John B. Wallingford

**Affiliations:** Division of Biological Sciences, Department of Molecular Biology, Nagoya University Graduate School of Science, Nagoya, 464-8602, Japan.; Institute for Frontier Life and Medical Sciences, Kyoto University, Kyoto, 606-8507, Japan; Department of Molecular Biosciences, University of Texas at Austin, 78712, USA.

**Keywords:** Actomyosin oscillation, Convergent extension, Planar cell polarity, Cell intercalation, Prickle2, PCP

## Abstract

Oscillatory actomyosin flows play a key role in single cell migration and in collective cell movements that shape invertebrates embryos, but the role of such oscillations in vertebrate morphogenesis remains poorly defined. Here, data from mathematical modeling and *in vivo* 4D imaging of actomyosin in the *Xenopus* gastrula suggest that oscillatory actomyosin contractions are a general feature of convergent extension by junction shrinking. We show that synchronous intracellular flows link two spatially distinct populations of actomyosin within individual cells, but that oscillations are asynchronous *between* neighboring cells that share a shrinking cell-cell junction. We also show that the core PCP protein Prickle2 displays a parallel oscillatory behavior and is required for tuning the frequency of actomyosin contractions, indicating that PCP signaling controls not only the orientation of actomyosin contractions, but also their frequency. Together, these data provide new insights into the function and control of oscillatory actomyosin contractions in collective cell movement.

## Introduction

Convergent extension (CE) is a ubiquitous process of collective cell movement that elongates the body axis in animals ranging from insects to mammals (Tada and Heisenberg, 2012). During convergent extension, cells rearrange by intercalating specifically in the mediolateral axis, thereby elongating the tissue in the perpendicular, anteroposterior axis (Walck-Shannon and Hardin, 2014). Two sub-cellular mechanisms have been shown to contribute to mediolateral cell intercalation (Shindo, 2017): First, mediolaterally-oriented protrusions act in a manner analogous to the leading edge of a migrating cell, driving cell intercalation via cell crawling (Shih and Keller, 1992). Second, pulsed actomyosin contractions actively shrink mediolaterally-oriented junctions (so called v-junctions, Figure S1A), thereby driving intercalation (Bertet et al., 2004, Blankenship et al., 2006). Recent work suggests that these two mechanisms act in concert in both vertebrates and *Drosophila* (Sun et al., 2017, Williams et al., 2014).

Actomyosin contraction is the key driver of animal cell movement (Gardel et al., 2010), so understanding how the actomyosin machinery is organized during CE is an important challenge. In the context of CE, this issue is currently best understood for junction shrinking in the epithelial cells of the *Drosophila* germband, a key model system for studies of cell intercalation (Bertet et al., 2004, Blankenship et al., 2006). In that case, two populations of actin have been shown to act in concert: Oscillation of a pool of “medial” actin drives pulsed constriction events at the junction, and oscillations of a second pool of “junctional” actin stabilizes the newly-shortened junction (Fernandez-Gonzalez and Zallen, 2011, Rauzi et al., 2010, Sawyer et al., 2011).

Cell intercalation has also been extensively studied in the mesenchymal cells of the *Xenopus* gastrula mesoderm (Figure S1B, Keller et al., 2000), though how actomyosin dynamics relate to cell intercalation in these cells is far less clear. As in the *Drosophila* germband, two actin populations have been described in the *Xenopus* mesoderm during CE. First, there is a “node-and-cable” system of actin filaments that resides superficially (i.e. 1–2 microns deep; Fig. S2A, A’) and displays oscillations that are implicated in cell intercalation by cell crawling (Kim and Davidson, 2011, Pfister et al., 2016, Skoglund et al., 2008). In addition, we have described a pulsatile actin population associated with junction shrinkage deeper within the tissue (4-6 microns; Fig. S2B, B’)(Shindo and Wallingford, 2014). Here, we will refer to these as the “superficial” and “deep” actin populations, respectively (see Figure S2). The relationship between these two actomyosin populations remains unclear.

Understanding actomyosin dynamics during CE in a vertebrate is crucial, because Planar Cell Polarity (PCP) signaling is an essential regulator of cell intercalation specifically in vertebrates (Butler and Wallingford, 2017) and not in *Drosophila* (Zallen and Wieschaus, 2004). Moreover, CE is essential for normal neural tube closure in vertebrates and PCP genes are key genetic risk factors for human neural tube closure defects (Wallingford et al., 2013). PCP proteins control myosin phosphorylation and disruption of PCP signaling severely disrupts myosin-mediated junction shrinking in diverse tissues, including the *Xenopus* mesoderm (Butler and Wallingford, 2017). However, despite the wide-spread implication of actomyosin oscillation and flow in morphogenesis of invertebrates (He et al., 2010, Fernandez-Gonzalez and Zallen, 2011, Martin et al., 2009, Rauzi et al., 2010, Sawyer et al., 2011) and in migration of single vertebrate cells in culture (Huang et al., 2013, Maiuri et al., 2015), we understand little about the spatial and temporal dynamics of actomyosin and how these are influenced by PCP proteins during vertebrate collective cell movement.

In this study, we adopted a mathematical modeling approach to ask how patterns of actomyosin behavior may impact cell intercalation during CE, and we performed high-resolution, three-dimensional time-lapse imaging of actin and myosin dynamics during CE in the dorsal gastrula mesoderm of *Xenopus*. Together with the previous findings in *Drosophila*, our findings *in silico* and in *Xenopus* gastrulae *in vivo* argue that oscillatory actomyosin contraction is a general principle of cell intercalation via junction shrinking that is conserved across cell types and animals. We also show that in addition to its known role in controlling the planar orientation of actomyosin contraction, PCP signaling is also required to tune the frequency of those oscillatory contractions. These results shed light on the molecular mechanisms of CE and of congenital diseases related to defective PCP signaling.

## Results

### Mathematical modeling suggests that asynchronous contractile oscillations with an optimal frequency increase the efficiency of convergent extension

Mathematical modeling has become a powerful tool for exploring complex biological events, including collective cell movements during morphogenesis (Szabó et al., 2016, Inoue et al., 2016, Alt et al., 2017, Yu and Fernandez-Gonzalez, 2017). Collective cell movements emerge *in vivo* from the coordination of actomyosin contractile systems both within cells and between cells (Guillot and Lecuit, 2013, Heisenberg and Bellaiche, 2013, Mao and Baum, 2015). Such multi-layered regulation makes understanding the significance of actomyosin dynamics challenging. We therefore developed a mathematical model to simulate the interplay of actomyosin contraction and junction shrinking during CE.

To this end, we modified the 2D vertex model, which is a well-defined model for simulating cell rearrangements and tissue deformation (Farhadifar et al., 2007, Staple et al., 2010). In the 2D vertex model, cell-cell junctions are modeled as two vertices and one edge shared by neighboring cells, and junction shrinking is modeled by tension exerted along edges thereby reducing the distance between vertices. We limited contraction to edges aligned along the x axis, representing the mediolateral contraction of cell-cell junction, as we previously observed in *Xenopus* (Shindo and Wallingford, 2014)(v-junctions, Figures 1A, 1A’, and S1A). We quantified the efficiency of tissue deformation by counting the number of new edges emerging after complete shrinking of a v-junction (Figure 1B). We then tested the effect of continuous versus oscillating contractions, and found that cells completely failed to intercalate when contraction of v-junctions was continuous and simultaneous (Figures 1C, 1G (blue), and S3A, Movie S1). By contrast, oscillating contractions did drive cell intercalation (Figures 1D, 1G (black), and S3B, Movie S1).

**Figure 1.**
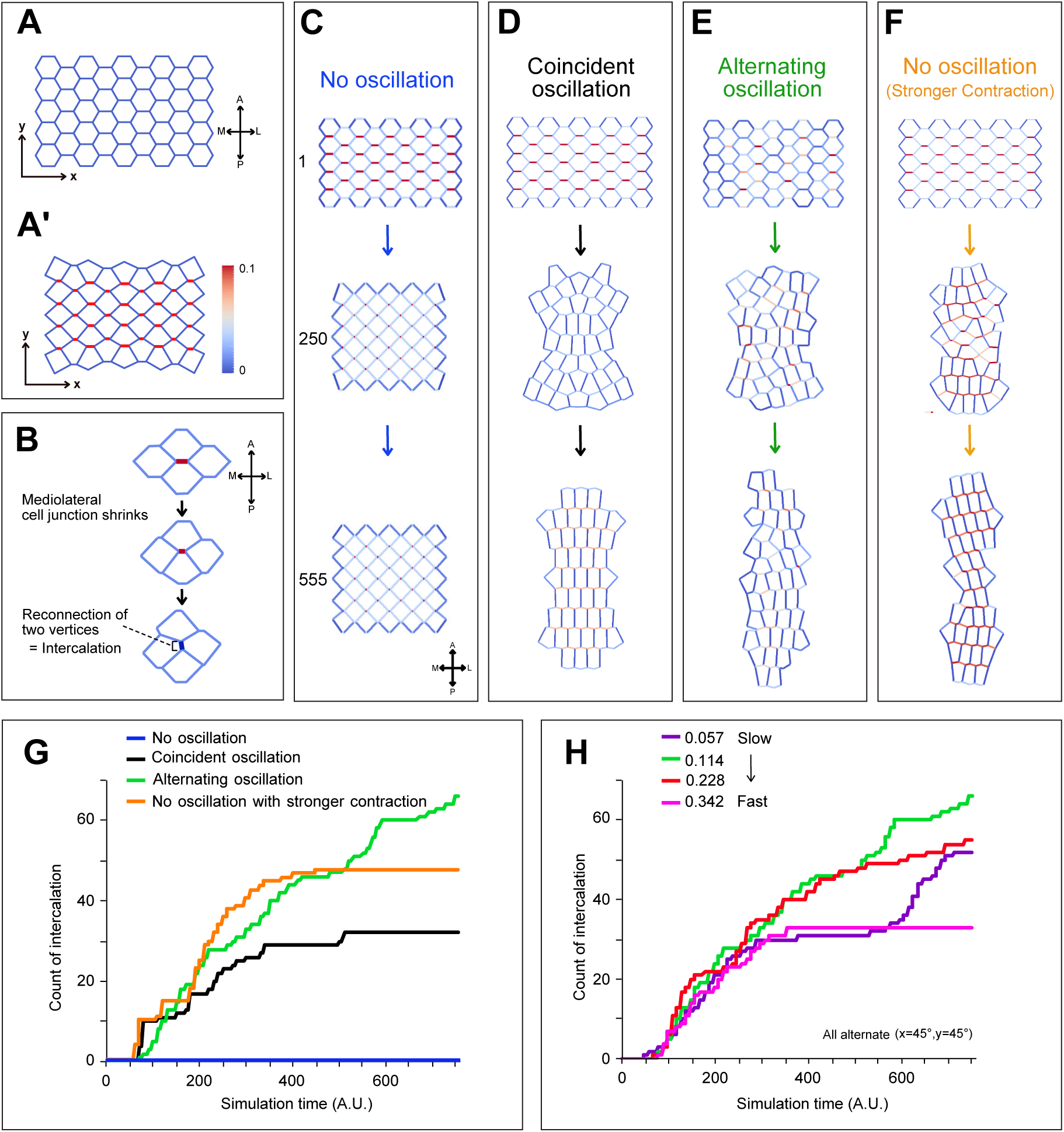
A 2D vertex model reveals that the alternate, carefully timed contractions are a general feature of cell intercalation. **(A)** Initial cell arrangement used in the simulations. **(A’)** Example of junction contractions. The cell-cell junctions aligned along the mediolateral axis (x axis) can contract in the model. The edge color indicates the magnitude of the line tension of the cell and cell junction (red=maximum strength, blue=minimum strength). **(B)** Example of the junction remodeling in the simulations. After the cell-cell junction shows emergence of a new junction, and is counted as one intercalation, as shown in Figures 1G and 1H. **(C)** Simulation of CE with coincident, continuous contraction without oscillation generates no intercalation (red indicates contraction). Top panel shows cells at 1 simulation time t = 1 (A.U); middle panels at t = 250 and lower panels at t = 555. Vertical axis indicates antero (A) - posterior (P), and horizontal axis indicates medio (M) – Lateral (L). **(D)** Coincident oscillating contractions (In-phase oscillation) generate modest intercalation. **(E)** Alternate oscillating contractions generate robust intercalation. **(F)** Coincident and continuous contraction without oscillating at 1.5 times the line tension generates modest intercalation. **(G)** Graph of intercalation quantified by the number of newly formed cell-cell junctions after complete v-junction shortening, representing the efficiency of cell intercalation. Blue: continuous contraction (Λ = 0.10); Black: In-phase oscillation (Λ = 0.10); Green: alternating oscillation (Λ = 0.10); Orange: continuous contraction (Λ = 0.15). Frequency of oscillation is defined by angular frequency *ω* [radian/simulation time unit] (*ω* = *2π*/*T*, See experimental procedures). For both in-phase and alternating oscillations, *2π*/*T* is fixed to 0.114. The phase shift to generate alternating oscillation is defined by θ_x_=0.7854, θ_y_=0.7854 (see experimental procedure). **(H)** Effect of oscillating frequency on intercalation efficiency for alternating oscillation. Purple: *ω* = 0.057 (slowest), Green: *ω* = 0.114, Red: *ω* = 0.228, Pink: *ω* = 0.342 (Fastest). The phase shift is fixed to Δθ = 2.5π×10^−1^ for the three groups.

Next, we explored the role of contraction timing by comparing the effect of coincident oscillations (i.e. all cells oscillate in phase, Figures 1D and S3B) to that of alternating oscillations. Strikingly, alternating oscillations between neighboring cells along anteroposterior axis resulted in significantly more cell intercalation than did coincident contractions (Figures 1D, 1E, and S3C, Movie S1). Moreover, with coincident oscillations, the rate of cell intercalation eventually stalled, reaching a stable plateau beyond which no additional intercalation occurred (Figure 1G, black). By contrast, when oscillations were alternating, the rate of intercalation remained roughly constant, and cell intercalations continued for the entirety of the simulation (Figure 1G, green). To probe the significance of this finding, we explored the interplay of oscillation timing and contraction strength. Strikingly, increasing the force of each contraction improved intercalation even in the absence of oscillations (Figure 1F, Movie S1), but importantly, it did not overcome the eventual plateau beyond which no more intercalation occurred (Figure 1G, orange). Thus, our model suggests that alternating contraction between neighboring cells that share a shrinking junction is an important factor for convergent extension.

Finally, our model provided us with the means to test the significance of contraction frequency in CE. We found that when contraction oscillation was very rapid, intercalation either slowed or plateaued (Figure 1H, red and pink). Decreasing contraction frequency dramatically improved intercalation (Fig. 1H, green), however only up to point; at very slow contractions frequencies, the progress of intercalation became irregular, showing a double plateau (Figure 1H, purple). Taken together, our simulations suggest that asynchronous and alternating oscillation, as well as tight control of contraction frequency, are important and general features that contribute to convergent extension by junction shrinking.

### Synchronized oscillation links intracellular actomyosin populations during convergent extension of the *Xenopus* mesoderm

To explore actomyosin oscillation in a vertebrate *in vivo*, we focused on the dorsal mesoderm of the frog, *Xenopus*. This animal is a well-established model for studies of CE (Keller et al., 2000, Shindo, 2017), as the large cell size allows for imaging of subcellular behaviors and protein dynamics with exceptional resolution (Kieserman et al., 2010). To provide additional resolution of actin behaviors in single cells within the collective, we performed mosaic labeling and imaged actin dynamics using three-dimensional time-lapse imaging of the dorsal gastrula mesoderm (“Keller” explants) (Figures S1C, D, D’).

We first asked if the node and cable actin system previously described to reside superficially in these cells (Figure S2A)(Kim and Davidson, 2011, Skoglund et al., 2008) is related to the actin population observed at cell-cell junctions deeper in the cell (Figure S2B)(Shindo and Wallingford, 2014). Using 3D time-lapse imaging, we observed that the superficial and deep actin pools oscillate synchronously in individual cells during junction shrinking (Figure 2A, Movie S2). To quantify this oscillatory behavior, we identified actin flows as the peaks of normalized fluorescent intensity over time (Figure S4), and this analysis revealed that actin pulses initiated in the deep population at shrinking cell-cell junctions and then flowed to the superficial actin pool (Figure 2B). Cross-correlation of the data confirmed the synchrony of deep and superficial actin pulses, with the peak of superficial actin intensity following the peak of deep actin by ~20 seconds on average (Figure 2C, max of cross-correlation coefficient: 0.404 ± 0.154 (mean ± sd), each cross-correlation: p < 0.001).

**Figure 2.**
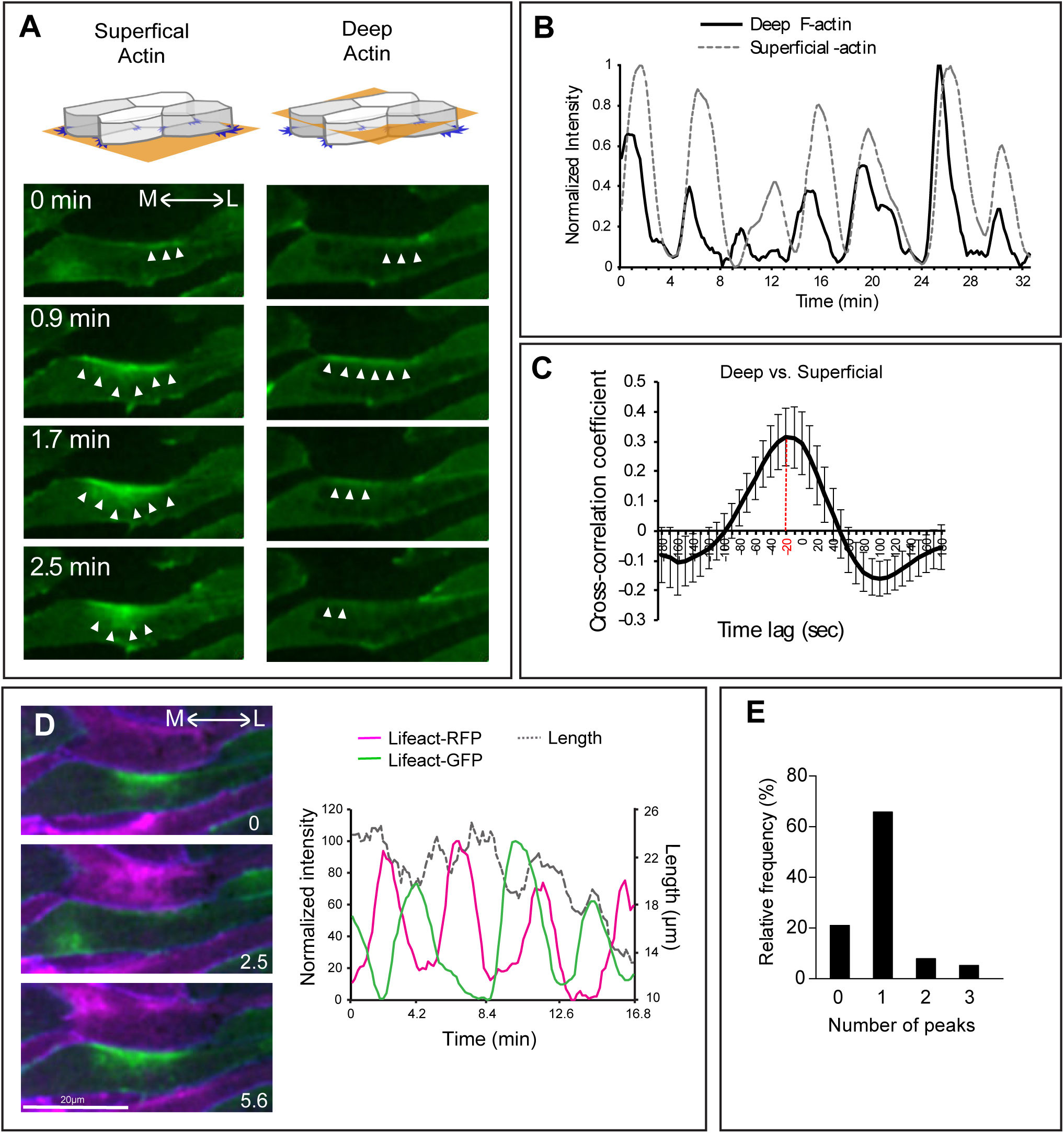
Actomyosin oscillations during convergent extension of the *Xenopus* dorsal mesoderm. **(A)** Mosaic expression of Lifeact-GFP allows imaging of labeled cells apposed to unlabeled cells for unambiguous assessment of actin dynamics in the superficial (left) and deep (right) actin populations. Stills from time-lapse movies show the progression of actomyosin flow across time and space (arrowheads). M: medial, L: lateral. **(B)** Peaks of normalized actin intensity allow quantification of actomyosin dynamics in the deep and superficial populations along contracting v-junctions from a single cell. **(C)** Actomyosin oscillations in deep and superficial populations are synchronous, showing a significant cross-correlation. **(D)** Two-color labeling of F-actin in same cells in panel A reveals that oscillations along shared v-junctions in apposed cells are out of phase; quantification of normalized intensity together with junction length over time is displayed at right. **(E)** The number of Lifeact’s peaks observed in a given cell between any two peaks occurring in an adjacent cell shows pulses in neighboring cell display a roughly one-to-one ration (See Figure S7).

In the *Drosophila* germband, actin oscillations are coordinated with myosin, and we previously reported that the myosin regulatory light chain Myl9 oscillates with the deep pool of junctional F-actin at shrinking-junctions during CE (Shindo and Wallingford, 2014). When co-expressed with Lifeact-RFP and membrane-BFP, we found that Myl9-GFP also oscillated with F-actin in the superficial population (Figure S5A, Movie S3). Cross-correlation analysis between the intensity of Myl9-GFP and Lifeact-RFP indicated a synchronized oscillation of Myl9 and F-actin in these two populations (Figure S5A’, max of cross-correlation coefficient: 0.728766 ± 0.051(mean ± sd), each cross-correlation: p < 0.0001). Together with the data from *Drosophila* and our mathematical model, these results suggest that oscillatory actomyosin contractions are a general feature of CE that spans cell types and organisms.

### Anteroposterior neighbor cells display asynchronous intercellular actomyosin oscillations during convergent extension

Our data suggest that deep and superficial actomyosin flows are synchronized *within* individual cells during convergent extension, but our modeling suggests that coordination of flow *between* cells is also a critical factor. Because of the known relationship of anteroposterior neighbor cells in junction shrinking (Shindo and Wallingford, 2014), we used a two-color imaging approach to unambiguously assess actomyosin dynamics in neighboring cells (Figure S1D). Time-lapse imaging of these explants demonstrated that actin flows in anteroposterior neighbors displayed asynchronous oscillations (Figure 2D). These oscillations appeared anti-correlated in time-lapse movies (Figure 2D) and kymographs (Figure S6). Similar oscillations were also observed using Myl9-GFP (Figure S5B), but their timing was highly variable, so we were unable to detect clear cross-correlation on average (Figure S5B’). However, quantification of the relative timing of pulses (Figure S7) revealed a predominantly one-to-one ratio of oscillations between neighboring cells (Figure 2E), suggesting a predominantly alternating relationship between neighboring cells. Thus, oscillations are synchronous between the deep and superficial actomyosin populations *within* each cell, but oscillations *between* cells are asynchronous.

### Prickle displays both spatial and temporal patterns of localization to cell-cell junctions during CE

Both modeling and imaging data suggest that convergent extension requires the careful calibration of several parameters, including the frequency of contractile flows and their asynchronous coordination between neighboring cells. To explore the molecular control of these oscillations, we turned our attention to the PCP signaling network, which has been implicated in control of both deep and superficial actomyosin dynamics in the dorsal mesoderm (Kim and Davidson, 2011, Shindo and Wallingford, 2014). Recently, we showed that the core PCP protein Pk2 controls CE in neural epithelial cells, where it displays a tight temporal and spatial correlation with actomyosin at shrinking junctions (Butler and Wallingford, 2018). We therefore used mosaic expression to generate labeled cells that were directly apposed to unlabeled cells in the dorsal mesoderm (Figure 3A). We then quantified the normalized fluorescence intensity at both anterior and posterior cell faces (Figure 3B, red, black). This analysis revealed that Pk2-GFP localized predominantly to the anterior face of cells engaged in CE (Figures 3C–3E, Movie S4).

**Figure 3.**
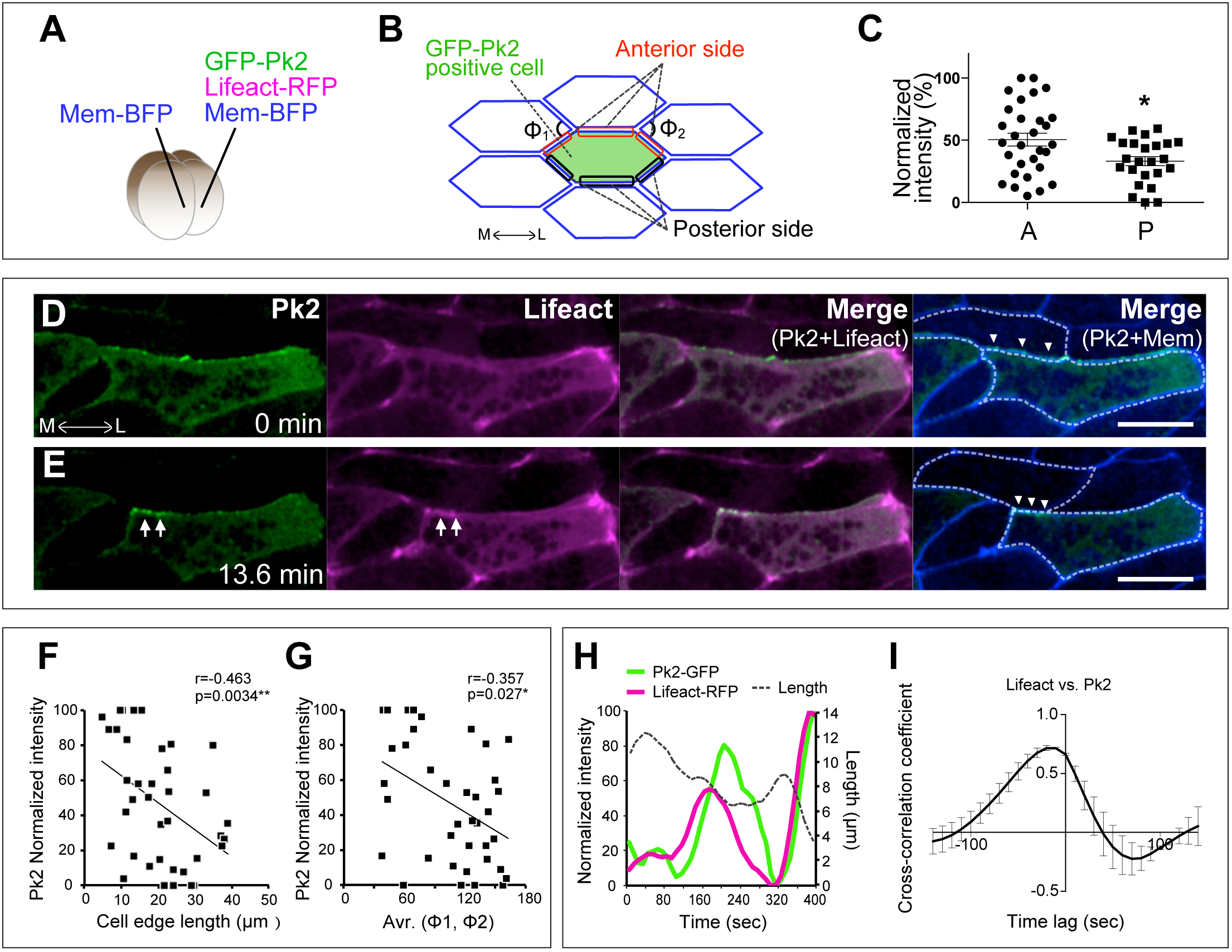
Prickle2 displays temporal and spatial patterns of localization during cell intercalation. **(A)** Method for mosaic expression of GFP-Pk2, Lifeact-RFP, and membrane-BFP in a *Xenopus* embryo at 4 cell stage. **(B)** Scheme showing the cell and cell-cell junctions measured for analysis in C, F, G, and H. The angle of apices of neighboring cells attached to a shrinking junction is indicated as Φ. **(C)** Mean intensity of GFP-Pk2 on anterior and posterior cell cortices. (* p = 0.0282, Student’s t-test; anterior: n = 30, posterior: n = 25) **(D, E)** Still images from time-lapse movie of GFP-Pk2 during junction shrinkage. Images of E are 13.6 minutes after the images in D. Arrowheads indicate contracting v-junction; arrows indicate the accumulation of GFP-Pk2 at the anterior face of the contracting v-junction. **(F)** Correlation between GFP-Pk2 intensity and the length of cell-cell junctions. GFP-Pk2 intensity is converted to percentages in each image. (38 junctions from 2 embryos). **(G)** Correlation between GFP-Pk2 intensity and the angle Φ. Same numbers as (F). **(H)** Quantification of normalized intensity of F-actin (Lifeact, magenta) and Pk2-GFP (green) together with junction length over time from single contracting cell junction. (I) Synchronous accumulation of F-actin and Pk2 is shown by cross correlation. Scale bar = 20μm.

While this spatial localization of Pk2 is consistent with data in other cell types and organisms (Ciruna et al., 2006, Jiang et al., 2005, Ossipova et al., 2015, Yin et al., 2008), our dynamic imaging allowed us also to detect interesting and novel *temporal* patterns. For example, we found that junctional Pk2-GFP intensity was elevated specifically at shrinking junctions (Figures 3D–3E). Indeed, increased junctional Pk2 intensity was significantly correlated with decreasing junction length (Figure 3F). Moreover, Pk2 intensity was also correlated with the angle ϕ (Figure 3B, 3G), a parameter that scales with the tension exerted on shrinking junctions during intercalation (Rauzi et al., 2010, Shindo and Wallingford, 2014). Further examination of Pk2 dynamics revealed a pulsatile pattern of junctional enrichment (Figure 3H), and we detected a significant cross-correlation between Pk2 intensity and actin intensity (Figure 3I, max of cross-correlation coefficient: 0.7155 ± 0.021 (mean ± sd), each cross-correlation: p < 0.0001). Thus, Pk2 displays both spatial and temporal patterns of localization during CE of the *Xenopus* dorsal mesoderm.

### Prickle2 controls the frequency of actomyosin oscillations

Finally, the coordinated pulsing of actomyosin and Pk2 is consistent with recent reports that indicate a reciprocal relationship in which actomyosin controls PCP protein localization and PCP proteins in turn control Myosin activity (Newman-Smith et al., 2015, Ossipova et al., 2015). Because we found that pulses of Pk2 were synchronized with those of actomyosin, we tested the effect of Pk2 loss of function on the several parameters of actomyosin oscillation defined above (i.e. Figures 1 and 2). We knocked down Pk2 using an antisense morpholino-oligonucleotide that has been validated in both convergent extension and polarization of multiciliated cells (Butler and Wallingford, 2018, Butler and Wallingford, 2015). Pk2 knockdown strongly disrupted both the elongation and the planar orientation of cells in the dorsal mesoderm, consistent with disruption of other core PCP proteins in dorsal gastrula mesoderm (Jessen et al., 2002, Wallingford et al., 2000)(Figures 4A–4D). Interestingly, the effect on actomyosin was strikingly specific, as Pk2 knockdown did not eliminate oscillations (Figures 4E–4H), did not alter the tight relationship between actin and myosin (Figures 4I, 4I’), and did not disrupt the asynchronous nature of oscillations between neighboring cells (Figures 4J, 4K, and S7, Movie S5). Instead, Pk2 knockdown specifically affected the frequency of actomyosin contractions, with knockdown cells displaying an inter-peak time roughly half that of controls (Figures 4F, 4H and 4J). Thus, our data suggest that Pk2 is required to tune the frequency of actomyosin oscillations, a parameter that our modeling suggests is essential for normal cell intercalation.

**Figure 4.**
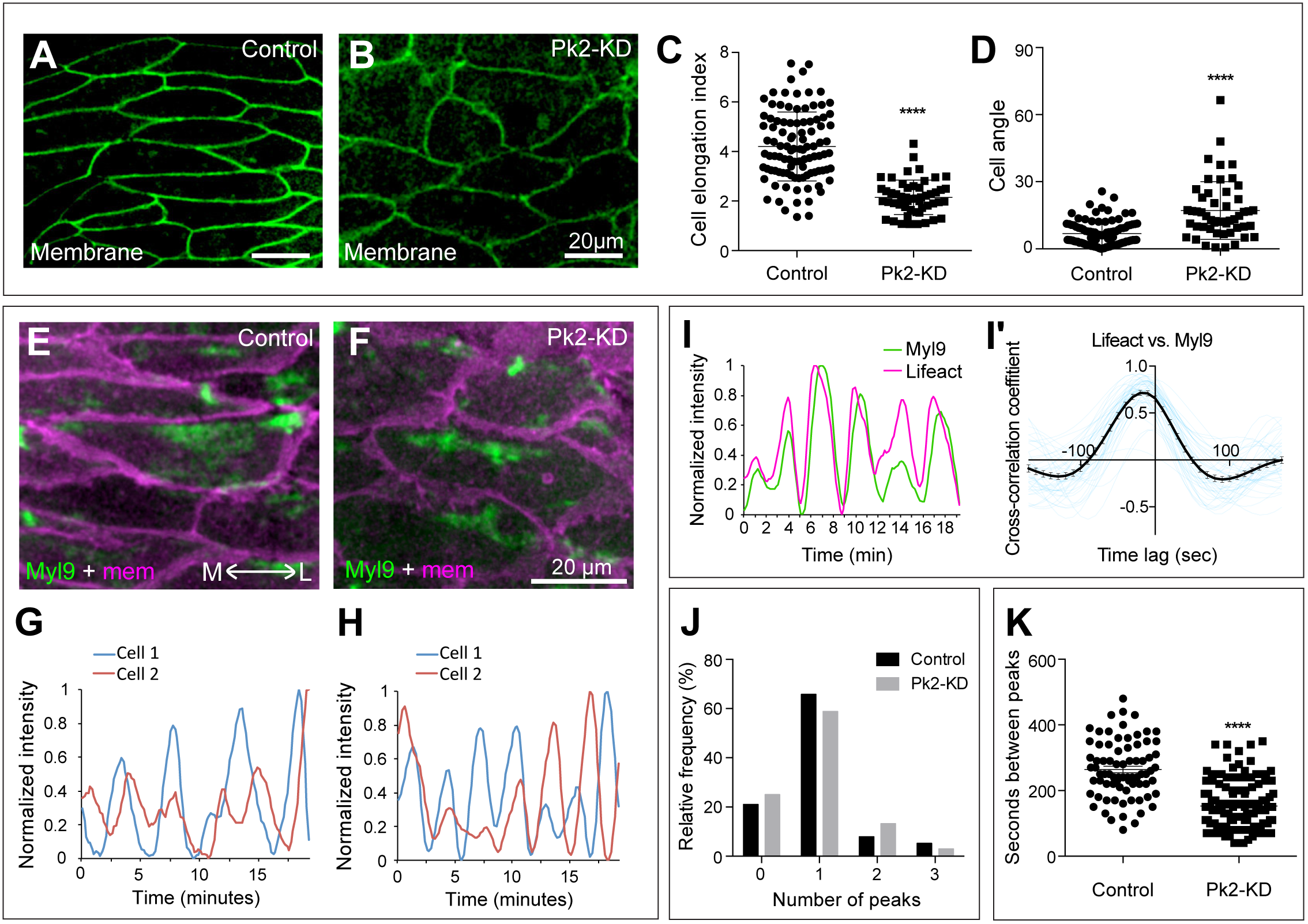
Prickle 2 controls the frequency of actomyosin pulses. **(A)** Cells in the *Xenopus* dorsal mesoderm display an elongate and aligned morphology. **(B)** Pk2 KD disrupts elongation and orientation of cells. **(C, D)** Quantification of cell shapes and orientations, respectively. (Cell elongation index: **** p < 0.0001, Student t-test, control: n = 100 from three embryos, Pk2-KD: n = 51 from three embryos; Orientation: **** p < 0.0001, Mann Whitney U-test, control: n = 97 from three embryos, Pk2-KD: n = 51 from three embryos). **(E, F)** Still images from time-lapse movies of Myl9-GFP and membrane-RFP in control (E) or Pk2-KD (F). **(G, H)** Representative oscillations of Myl9 intensity in two apposed cells in a control (G) or Pk2-KD (H). **(I)** Normalized intensities of Myl9-GFP and Lifeact-RFP, measured along mediolaterally aligned cell junctions in Pk2-KD embryos. **(I’)** Cross-correlation of normalized intensities of Myl9 and Lifeact along in Pk2-KD embryos revealed their synchronized oscillations (black line with SE). Each blue line is from each cell junction. **(J)** Peaks in in apposed cells generally display a one-to-one ration, suggesting a roughly asynchronous and alternate relationship between cell neighbors. See also Figure S7. **(K)** Time between peaks is significantly reduced after Pk2-KD. **** p < 0.0001, Mann Whitney U-test, control: n = 76 from three embryos, Pk2-KD: n = 138 from three embryos. Scale bar = 20 μm.

## Discussion

Oscillatory actomyosin contraction is a key feature of cell movement both in individual cells and in cell collectives, and such oscillations have been extensively defined during CE of the *Drosophila* germ band. However, CE is a ubiquitous morphogenetic process in animals, and we know little of such oscillatory contractions in other settings. Guided by mathematical modeling of cell intercalation by junction shrinking, we have quantified the patterns of actomyosin flow in the *Xenopus* dorsal mesoderm, a key model for studies of cell intercalation. The tightly cross-correlated flows observed between deep and superficial actin populations we observed within single cells are reminiscent of the flows between “medial” and “junctional” actin pools described in *Drosophila* (Rauzi et al., 2010, Sawyer et al., 2011). The asynchronous and alternate oscillations observed here between neighboring cells in *Xenopus* mesoderm are likewise reminiscent of similar patterns observed in *Drosophila* (Fernandez-Gonzalez and Zallen, 2011). Such similarities are highly significant because intercalations studied in *Drosophila* occur in tightly adherent epithelial cells with a well-defined apicobasal polarity, while the mesenchymal cells observed here in *Xenopus* are more loosely adherent and lack obvious apical-basal polarity. Moreover, CE in the *Xenopus* mesoderm is entirely dependent upon intact PCP signaling (Butler and Wallingford, 2017), but the analogous process in *Drosophila* is independent of this signaling system (Zallen and Wieschaus, 2004). Given these dramatic differences, the results of our modeling *in silico* (Figure 1) and imaging *in vivo* (Figure 2) strongly argue that asynchronous oscillatory contraction is a general principle of cell intercalation via junction shrinking that is conserved across cell types and animals.

Our data also provide important new insights into the function of PCP signaling in vertebrate CE, which is an important issue given the implication of PCP genes in human neural tube defects (Wallingford et al., 2013). These results here are consistent patterns we observed recently in neural epithelial cells (Butler and Wallingford, 2018), which again is significant for demonstrating that the dynamic spatial and temporal patterns of Pk2 localization are conserved across cell fate (ectoderm vs. mesoderm) and tissue type (epithelial and mesenchymal). These results are also consistent with findings that anteroposterior patterning is central to the organization of CE in both *Drosophila* and *Xenopus* (Paré et al., 2014, Ninomiya et al., 2004, Zallen and Wieschaus, 2004).

Finally, previous work has implicated PCP signaling in control of both the deep and the superficial actin populations in the *Xenopus* dorsal mesoderm (Kim and Davidson, 2011, Shindo and Wallingford, 2014), so our finding that actomyosin flow links these two populations is informative. In fact, disruption of the core PCP protein Dvl reduces the frequency of superficial actin contractions in these cells (Kim and Davidson, 2011), so our finding of the opposite effect following Pk2 loss of function (Figure 4) is consistent with the antagonistic relationship between these two proteins in *Drosophila* (Jenny et al., 2005, Tree et al., 2002). Perhaps most significantly, we show here that Pk2 displays an oscillatory pattern of junctional enrichment that correlated with that of actomyosin in mesenchymal cells of the dorsal mesoderm (Figure 3). Together with the observed enrichment of Pk2 specifically at shrinking junctions (Figure 3), these data suggest that the *temporal* patterns of PCP protein enrichment at cell-cell junction are at least as important as spatial patterns (i.e. planar asymmetry). These data highlight the essential role for dynamic analysis of PCP function in dynamic events such as CE and provide a foundation for future studies of the molecular control of actomyosin dynamics during vertebrate collective cell movement.

## Acknowledgements

Work in the A.S group is supported in part by funds from The Sumitomo Foundation, Tomizawa Jun-ichi & Keiko Fund of Molecular Biology Society of Japan for Young Scientists, and JSPS KAKENHI Grant Numbers JP15H01318, JP15K21065, JP26891012. Work in the M.K. lab is also supported by JSPS KAKENHI. Work in the Wallingford lab was supported by an R01 grant from the NIGMS and an R21 from the NICHD to JBW and a Uehara Fellowship to A.S. We thank Keita Ohsumi, Mariko Iwabuchi and Tomoko Nishiyama for sharing frog facility in Nagoya University.

## Author Contributions

A.S. designed and performed all *Xenopus* experiments, analyzed the results, and executed computational simulations. Y.I. designed, programed, and executed computational simulations. A.S. and J.B.W. conceived the project and wrote the manuscript. A.S., M.K. and J.B.W. oversaw the project.

## Experimental Procedures

### Preparation of *Xenopus* embryos

Ovulation of adult *Xenopus laevis* females was induced by an injection of human chorionic gonadotropin. After incubating at 16 °C overnight, eggs were squeezed from the females, fertilized *in vitro* and dejellied in 3% cysteine (pH 7.8–8.0) at the two-cell stage. Fertilized embryos were washed and subsequently reared in 1/3 x Marc’s Modified Ringer’s (MMR) solution. To prepare the embryos for mRNA microinjections, the embryos were placed in 1/3 x MMR solution with 3% ficoll.

### Plasmids and morpholino anti-sense oligonucleotides (MO)

GFP-Pk2 plasmids were constructed and designed as described previously in (Butler and Wallingford, 2015), and the Myl9-GFP plasmid in (Shindo and Wallingford, 2014). The Pk2 MO used disrupts splicing of the Pk2 gene (Butler and Wallingford, 2015) and severely disrupts CE of Keller explants and neural tube closure in intact *Xenopus* embryos; both phenotypes are rescued by co-expression of the wild-type Pk2 gene (Butler and Wallingford, 2018). Furthermore, these phenotypes are recapitulated by expression of a Pk2 dominant-negative (Butler and Wallingford, 2018) and are consistent with CE defects in a genetic mutant in the single *prickle* homolog in Ciona (Jiang et al., 2005).

### Morpholino and mRNA microinjections

Capped mRNAs for fluorescent markers (GFP-Pk2, Lifeact-GFP, Lifeact-RFP, Myl9-GFP membrane-RFP, membrane-BFP) were synthesized using a mMessage mMachine kit (Thermo Fisher Scientific, Ambion). Morpholino (MO) or mRNA was injected into the two dorsal blastomeres targeting the dorsal marginal zone (DMZ). Pk2-MO was injected at 30ng per blastomere. The amounts of injected mRNAs per blastomere were as follows: GFP-Pk2 [90 pg], Lifeact-GFP/RFP [60 pg], Membrane-RFP/BFP [60 pg], Myl9-GFP [20 pg].

### Live imaging of Keller explants

After mRNA injection, DMZ tissues were dissected out at stage 10.5 with forceps, hair loop, and heir knives (Figure S1C). The explants were mounted onto a fibronectin (Sigma, F1141) – coated glass bottom dish in DFA (Danilchik’s for Amy) medium, and cultured at 13 °C for overnight. The explant imaging with a confocal microscope (CV1000, Yokogawa Electric Corporation) was started when the sibling embryos reached at stage 14–15. The time-lapse movie was taken every 10 sec for recording actomyosin pulsing.

### Computational Simulation

We employ a 2D vertex model (Nagai T. and Honda 2001) to simulate the dynamics of multiple cells with cell rearrangement during convergent extension. In the model, cell shapes and their packing geometry are represented by a network of polygons consisting of vertices and edges. Cell rearrangement is expressed by implementing a local network topology change known as the T1 transition rule. The cell shape change is expressed by movements of vertices. The equation of motion of the *i*-th vertex at time *t* is

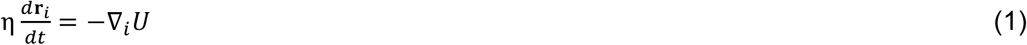

where η is a frictional coefficient, and **r**_*i*_ is the position vector of the *i*-th vertex. The mechanical properties of the cells are expressed by an energy function *U* as follows.

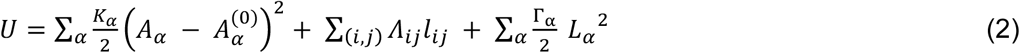

Here, the *α*-th cell’s area *A_α_*, perimeter *L_α_*, and junctional length *l*_ij_ between the *i*-th and *j*-th vertices are represented as variables. The constants *Κ*_*α*_ and Γ_*α*_ are the area elasticity and peripheral contractility, respectively. The energy function *U* in Eq. (2) is basically the same as suggested in (Farhadifar et al., 2007) (Staple et al., 2010), while we redefine the line tension *Λ_ij_* as a function of time and position of vertices to take into account a preferential direction of the oscillatory constriction at each individual cell described below.

The line tension Λ_*ij*_ on the junction line between *i*- and *j*-th vertices is decomposed to two contributions: orientation and oscillatory factors.

The orientation factor, *p_ij_*, is introduced to express the mediolateral-oriented constriction.

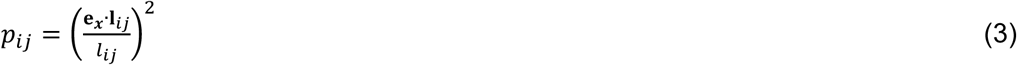

Here, e_*x*_ is the unit vector in the direction of the *x*-axis which is defined by the mediolateral axis (x axis in Figure 1A). The junctional line vector, **l**_*ij*_, is defined by the relative position vector from *i*-th to *j*-th vertices.

The oscillatory factor, Φ_*α*_(*t*) is introduced to express an oscillatory contribution from the *α*-th cell:

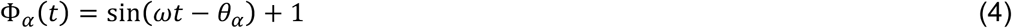

Here, the angular frequency, *ω* is identical for all cells where no oscillation occurs at *ω* = 0 (Figure S3B). The phase difference, *θ_α_*, is defined by the initial configuration (*n_x_, n_y_*), of the *α*-th cell:

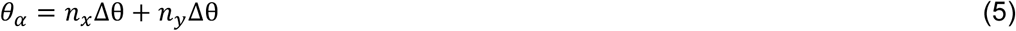

where, *n_x_* and *n_y_* are row and column indices for the *α*-th cell in the initial configuration of the cell packing, respectively. Δθ is a configuration-based phase shift where a coincident (in-phase) oscillation is reproduced by Δθ = 0 (Figure S3B).

Supposing that the junctional line between *i*-th and *j*-th vertices is shared by the *α*- and *α*\*-th cells, the line tension contributed from these two cells at the junction line is expressed as

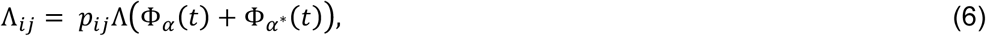

where Λ corresponds to a time-averaged value of the line tension. All model constants are listed in Table 1.

**Table 1.** Model parameters

| Symbol | Value | Descriptions |
| --- | --- | --- |
| $K_{\alpha}$ | 1.0 | Area elasticity of Eq. (2) |
| $\Gamma_{\alpha}$ | -6.0 | Peripheral contractility of Eq. (2) |
| $\Lambda$ | {0.10, 0.15} | Mean value of line tension of Eq. (6) |
| $A_{\alpha}^{(0)}$ | 2.6 | Cell area at stress free state of Eq. (2) |
| $\omega$ | {0, 0.57, 1.14, 2.28, 3.42}$\times 10^{-1}$ | Angular frequency of Eq. (4) |
| $\Delta\theta$ | {0.0, $2.5\pi \times 10^{-1}$} | Configuration-based phase shift of Eq. (5) |
| $\eta$ | 1.0 | Friction coefficient of Eq. (1) |
| $\Delta t$ | $1.0 \times 10^{-1}$ | Time step size for numerical integration of Eq. (1) |

### Image and statistic analysis

Images were quantified by using Fiji (https://fiji.sc/) and Igor Pro. Statistic analyses were performed using Prism6, GraphPad Software, Inc. The statistic test for each analysis is described in the figure legends. Cross-correlation coefficient was calculated using R (https://www.r-project.org/).

## Supplemental Information

**Figure S1.**
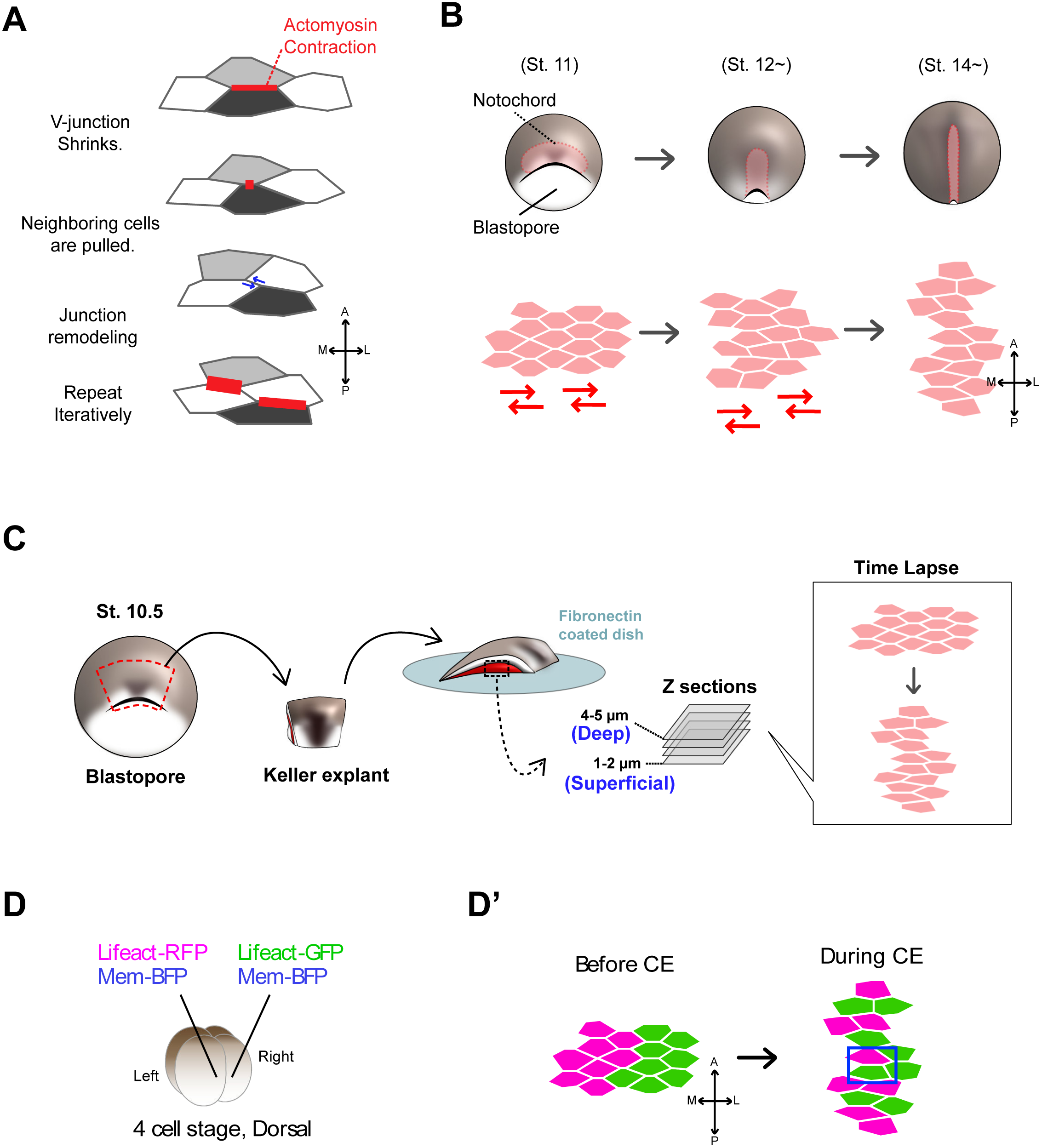
Cell movements during convergent extension in *Xenopus* notochord. **(A)** Driving force of cell intercalation along medio-lateral axis. Myosin light chain is phosphorylated along the cell-cell junction aligning mediolateral axis (red line, so called v-junction), and contracts the cell junction. The neighboring cells (white colored cells) are pulled by the junction contraction, then sequentially cause junctional remodeling (blue arrows). After such cell rearrangement, newly formed v-junction contracts by the actomyosin. **(B)** Notochord is formed during the process of gastrulation to neurulation at dorsal midline (St. 10.5–St. 18). The notochord (pink color) elongates along anteroom-posterior axis as blastopore closes by CE. The cells in notochord elongate and move along medio-lateral axis to intercalate (cells move along the red arrows). **(C)** Keller explant is isolated from dorsal mesoderm at St. 10.5 *Xenopus* embryos for live-imaging of CE. The notochord cells in Keller explant undergo normal CE as observed in an intact whole embryo. Live imaging of the cell movement is taken by mounting the Keller explant on a glass bottom dish coated with fibronectin, and monitored with an inverted confocal microscope. The Z-plane of images in Figure 1A is around 1–2 μm from the glass (superficial), and of images in Figure 1B is taken around 4–5 μm from the glass (deep). **(D)** Mosaic expression of Lifeact-RFP, Lifeact-GFP, and membrane-BFP in a 4-cell stage of *Xenopus* embryo. By undergoing CE, the cell populations labeled with different colors are mixed, allowing us to recognize F-actin at the cell-cell junction in both cells sharing the junction.

**Figure S2.**
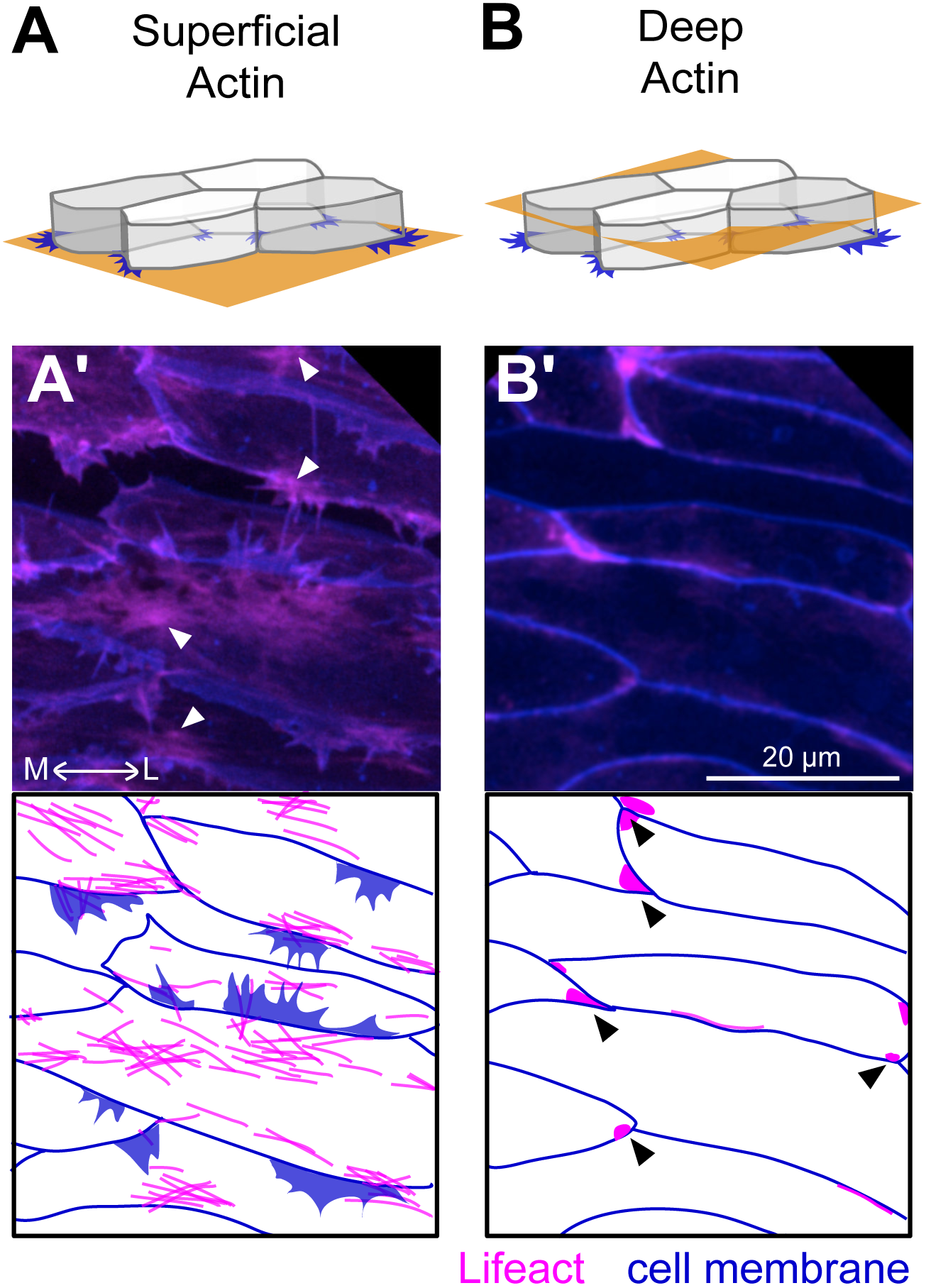
Two distinct Z-planes observed in *Xenopus* notochord cells to detect medial or junctional actin and myosin. **(A, B)** Schemes of notochord cells in an isolated Keller explants. Yellow plane indicates the observed Z-plane, a superficial plane for medial actomyosin (A), and a deep plane for junctional actomyosin (B), respectively. **(A’, B’)** Fluorescent images of notochord cells expressing Lifeact-RFP and membrane-BFP in the superficial (A’) and deep (B’) Z-plane. Node-and-cable is visible as medial actin in the superficial plane, while junctional actin is visible more in the deep plane. White arrowheads in A’ indicate “node”, and black arrowheads in B’ indicate F-actin accumulations at the multicellular junctions.

**Figure S3.**
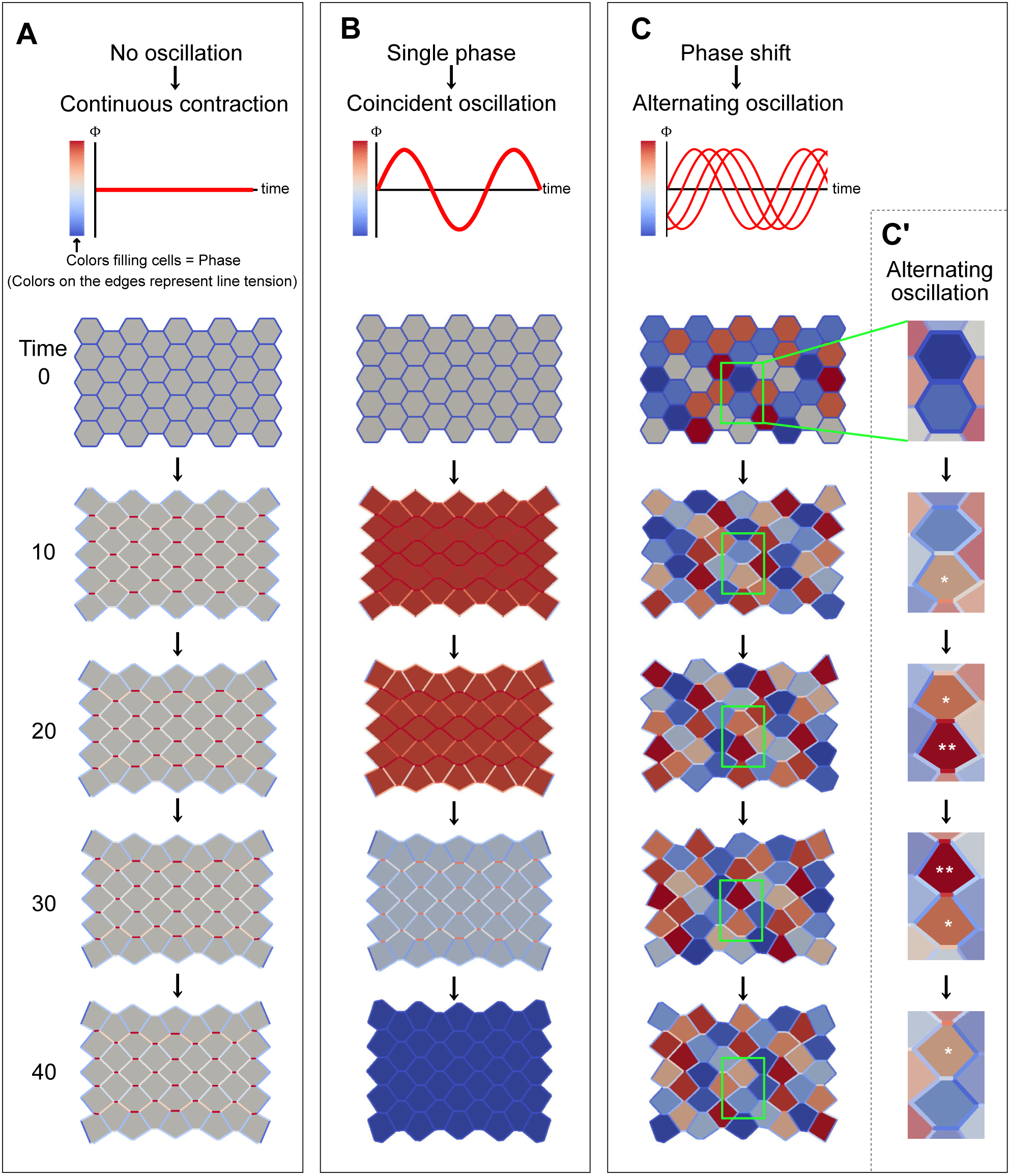
Designing 2D vertex model for comparison of different contraction modes. The graphs indicate change of oscillatory factor, Φ (Eq. 4), with respect to time in each mode. Color bar in left side of the graph is to indicate value of oscillatory factor in each cell, representing a temporal magnitude of line tension exerted by each cell. Note that the colors on the edges in the simulations show the resultant line tension exerted by neighboring two cells (Eq. 6), as shown in Figure 1A. **(A)** Continuous contraction without oscillation at angular frequency ω= 0 and Δθ = 0, which represents no spatiotemporal phase change as shown by gray color filling all cells uniformly. **(B)** Coincident oscillation exerted by single phase, and defined by Δθ = 0. All cell contract simultaneously, represented by uniform color changes through all cells. **(C)** Alternate oscillation exerted by phase shift. Each cell contracts in different but coupled timing, which can generate alternate oscillation along anteroposterior axis. **(C’)** Example of alternating oscillation in the simulation. The cell providing force to shorten sharing edge is alternating, that can be recognized by the cell colors. Asterisk and its number indicate the cell exerting force and its magnitude to shorten mediolateral edge, respectively. The phases between the two cells are coupled, representing alternating oscillations.

**Figure S4.**
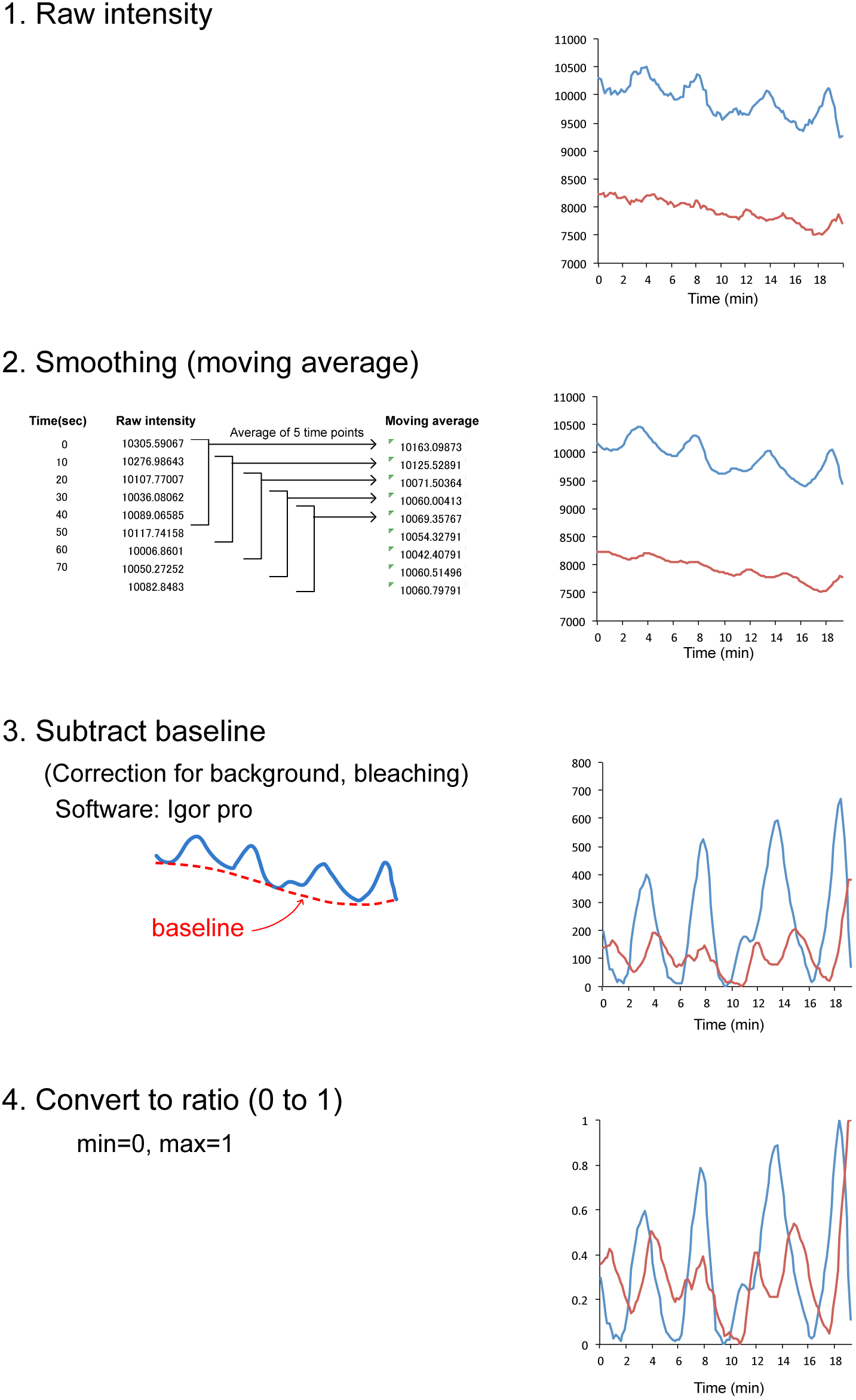
Normalization of measured intensities of oscillated actin and myosin. Raw intensities of target GFP or RFP were measured along contracting v-junction (1). Moving average was calculated to smooth each line (2). To avoid effects from bleaching during taking time-lapse, baseline was subtracted using IgorPro (3). The intensities were converted to ratio (4). Note that the normalization is to compare timing of accumulations of the target proteins along single contracting v-junction, and not to compare absolute amount of each proteins.

**Figure S5.**
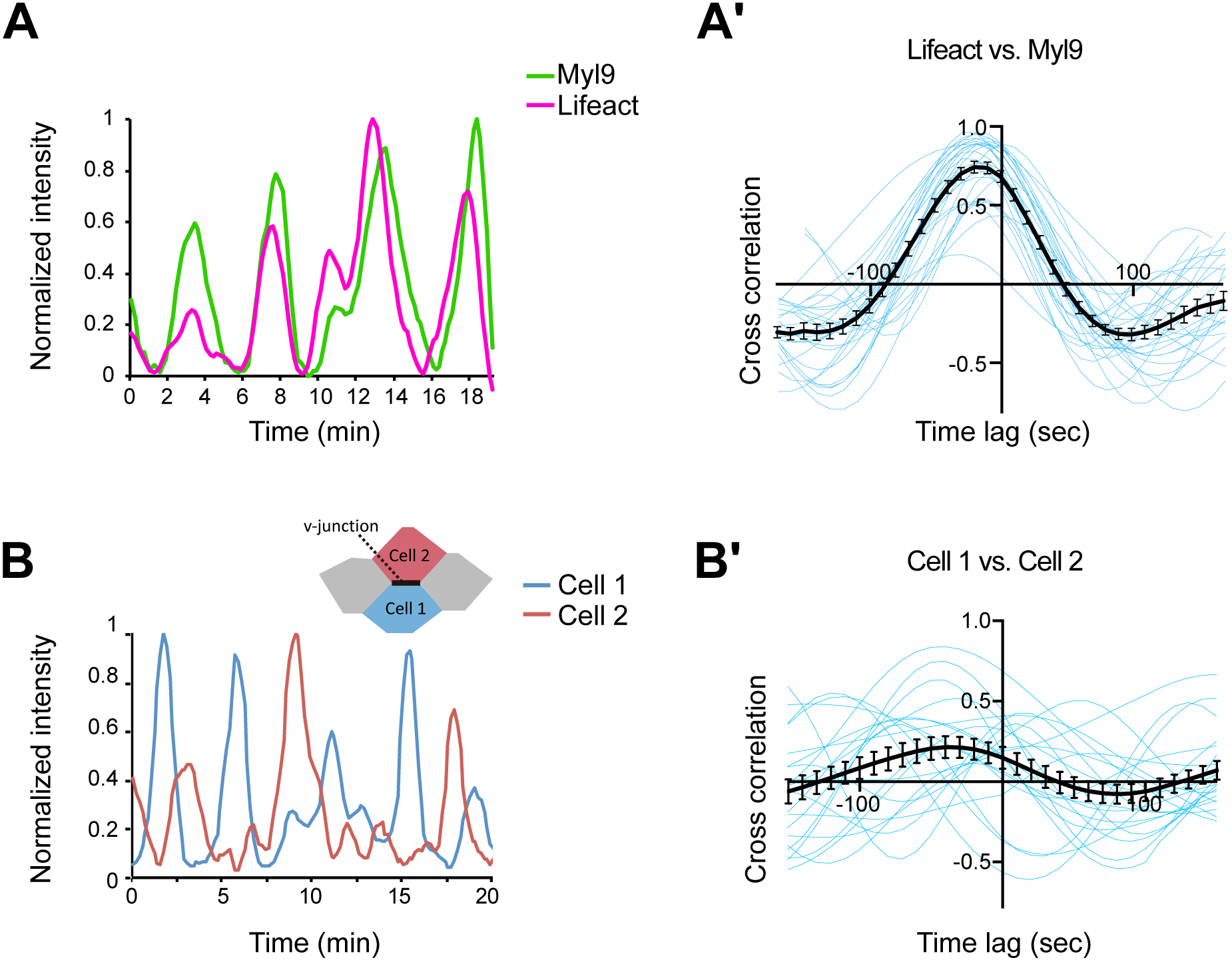
Myl9 oscillations coupled with F-actin along contracting v-junctions. (A) Normalized intensities of Myl9-GFP and Lifeact-RFP, measured along contracting v-junction. (A’) Cross-correlation of normalized intensities of Myl9 and Lifeact along contracting v-junction revealed their synchronized oscillations (black line with SE). Each blue line is from each v-junction. (B) Normalized intensity of Myl9-GFP in adjacent cells composing contracting v-junction. (B’) Cross-correlation of normalized intensities of Myl9-GFP in adjacent cells composing contracting v-junction (black line with SE). Each blue line is from each v-junction, showing various time lags.

**Figure S6.**
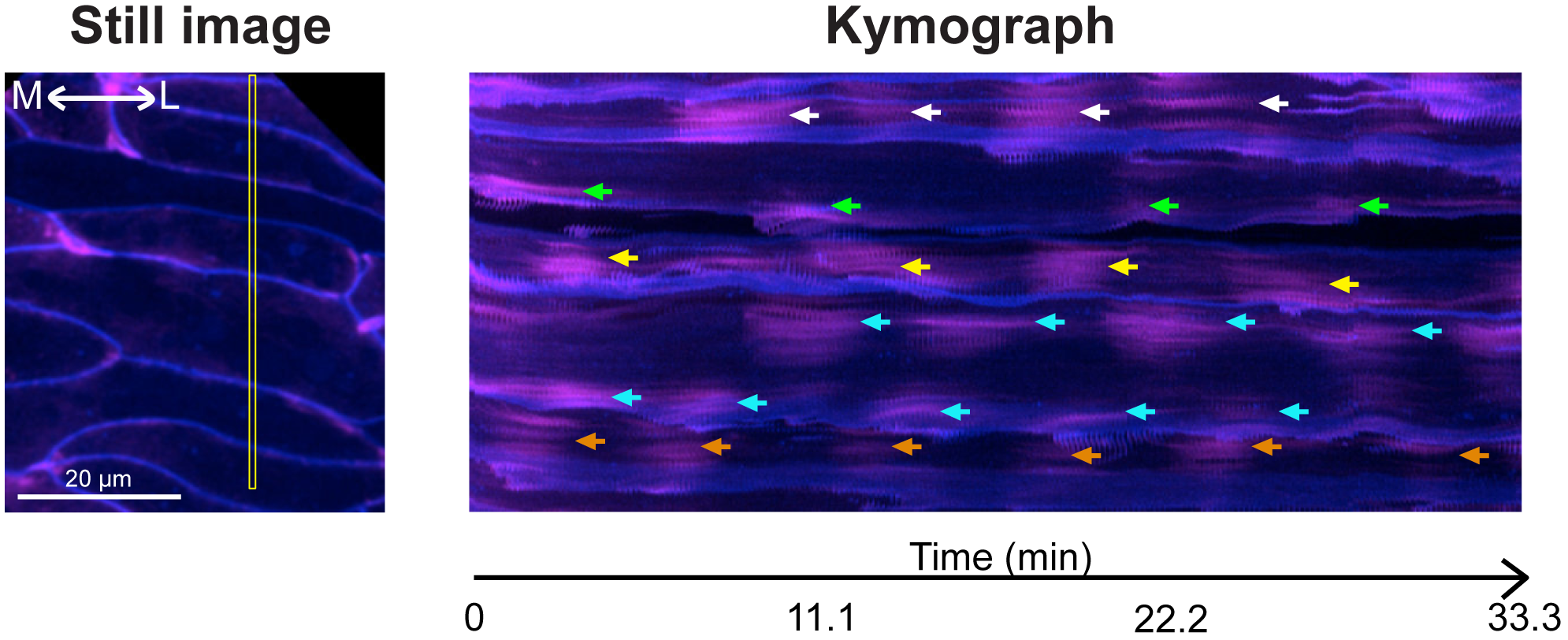
Cortical actin along mediolateral cell-cell junctions display asynchronous oscillations. Kymograph of superficial plane generated from time-lapse movie of Lifeact-RFP and membrane-BFP shown in Figure S2. Yellow box in the left panel indicates the region detected for making kymograph. Each color of arrows shows F-actin accumulations along mediolateral cell-cell junctions in each cell.

**Figure S7.**
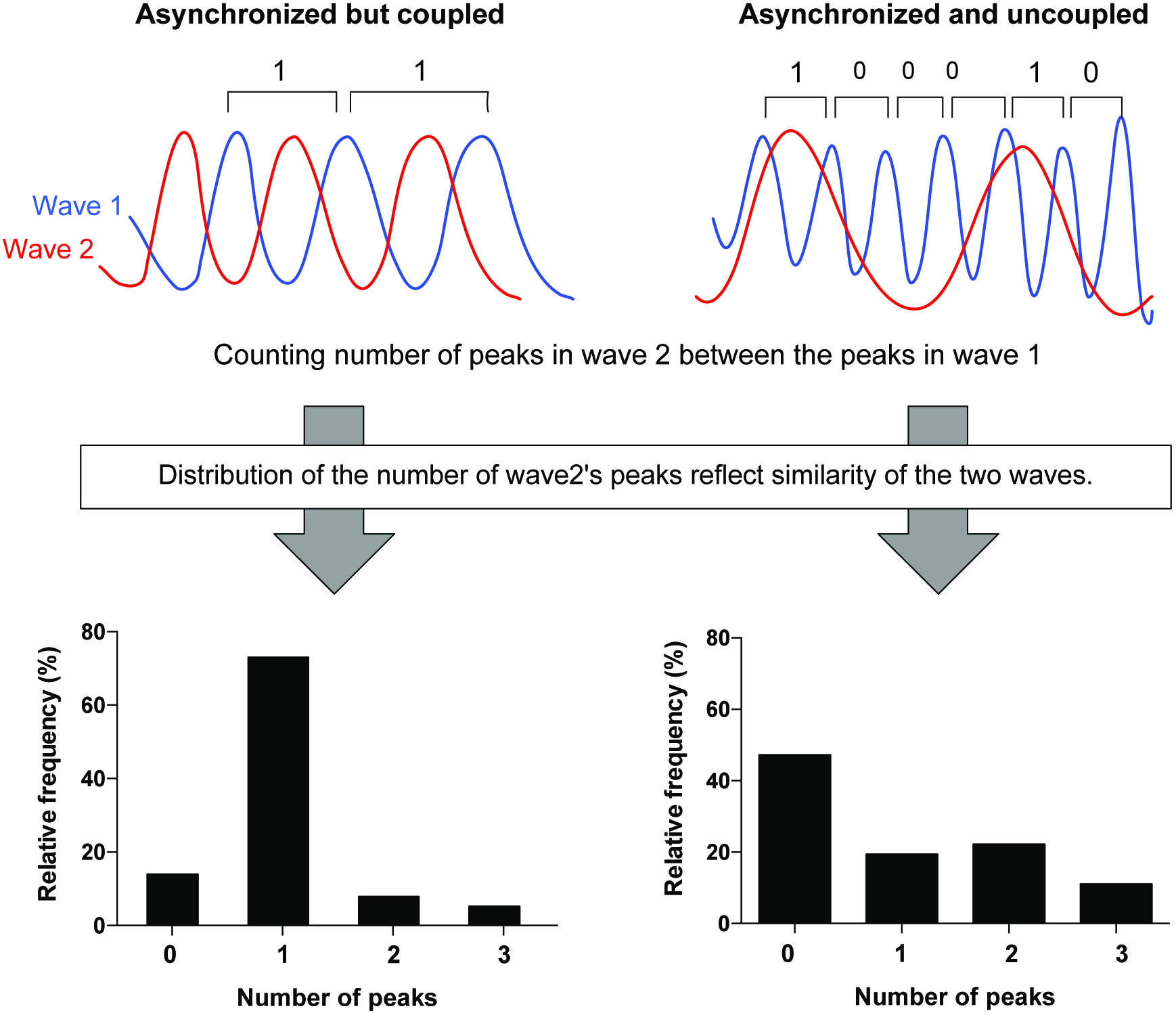
Quantification of alternating, asynchronous oscillations. Relative timing of pulses was quantified by counting the number of peaks in the wave (red) between the peaks in another wave (blue). If two waves are asynchronous but alternating as shown in left side, the number converges on 1. If two waves are asynchronous and unrelated as shown in right side, the number has a variety.

**Movie S1**

Simulation of CE with different timing of actomyosin contraction. **(A)** Coincident and continuous contraction without oscillation (Λ=0.1). **(B)** Coincident contraction with oscillation (Λ=0.1). **(C)** Alternating oscillation (Λ=0.1). **(D)** Coincident and continuous contraction without oscillation, but higher line tension (Λ=0.15).

**Movie S2**

Medial or Junctional F-actin (green) co-expressed with membrane-BFP (magenta) during v-junction constriction. Left two panels are superficial (medial) actin, right two panels are deep (junctional) actin. Bar = 10 μm. 12 frames / sec.

**Movie S3**

Asynchronous oscillations of medial F-actin (magenta) and Myl9 (green) with membrane-BFP (blue). From left side, merging of Myl9 and cell membrane, F-actin and cell membrane, and all three colors. Bar = 20 μm. 12 frames / sec.

**Movie S4**

Prickle accumulation at the shrinking v-junction. Pk2-GFP (green), Lifeact-RFP (magenta), and membrane-BFP (blue). From left side, merging of Pk2 and cell membrane, Pk2 and F-actin, and all three colors. Bar = 20 μm. 12 frames / sec.

**Movie S5**

Medial Myl9 pulsing (green) in control or Pk2-KD embryo. Membrane is labeled with BFP (blue). Bar = 20 μm. 12 frames / sec.

